# Population analysis and host-disease associations of Shiga toxin-producing *Escherichia coli* from various sources across eleven European countries using whole genome sequencing

**DOI:** 10.64898/2026.04.27.721056

**Authors:** Rosangela Tozzoli, Tristan Schadron, Arnold Knijn, Luca De Sabato, Stefano Morabito, Margherita Montalbano di Filippo, Eve Fiskebeck, Gro S. Johannessen, Jeevan Karloss Antony-Samy, Linnea Good, Robert Söderlund, Angela H. A. M. van Hoek, Lapo Mughini-Gras, Eelco Franz, Kinga Wieczorek, Gaia Scavia, Ornella Moro, Paola Chiani, Valeria Michelacci, Catherine Burgess, Geraldine Duffy, John Rodgers, Miranda Kirchner, Angela Pista, Leonor Silveira, Ana Amaro, Lurdes Clemente, Marie A. Chattaway, Claire Jenkins, Timothy J. Dallman, Susanne Schjørring, Flemming Scheutz, Brian Byrne, Montserrat Gutierrez, Vicente Lopez-Chavarrias, María Ugarte-Ruiz, Lin Brandal, Umaer Naseer, Ivana Kolackova, Aldert L. Zomer, Jaap A. Wagenaar, Sara M. Pires, Tine Hald, Camilla Sekse

## Abstract

Shiga toxin-producing Escherichia coli (STEC) are important foodborne pathogens, able to cause severe disease in humans. In the *DiSCoVeR* project (https://onehealthejp.eu/jrp-discover/) a STEC inventory from human and non-human sources from 11 European countries was set up and ≥ 3500 strains were sequenced to perform comparative genomics analysis. We used this dataset to assess STEC population structure and to investigate potential associations between genomic features, host reservoirs and symptoms.

Most STEC isolates analysed by Whole Genome Sequencing (WGS) in this study were collected between years 2010-2020. An *ad hoc* pipeline was deployed for a harmonised characterization of the STEC in the database, allowing the determination of serotyping, *stx* gene subtyping, 7-loci MLST, virulotyping and cgMLST. The results were analysed with Principal Component Analysis (PCoA) in relation with isolation source to assess clustering of STEC subpopulations.

When human STEC data were analysed, the PCoA revealed three distinct human STEC subpopulations (STEC_1, STEC_2 and STEC_3), which were further analysed for associations between genomic features, symptoms and variance. The non-human STEC showed a more dispersed distribution, except for one subpopulation with genes linked to specific host species, and some virulence profiles overlapping with the STEC_1 population.

In conclusion, our analysis identified distinct STEC subpopulations from human cases, each characterized by specific genetic features and associated with varying proportions of severe disease outcomes. These findings provide novel insights supporting the risk assessment of STEC.

**Impact statement:** [*This lay summary of your article should be no more than 200 words, and should a) provide a perspective of how this article adds to the literature in the field; b) identify breadth of interest/utility; and c) state the significance of output (incremental or step), in terms of relevance*.]

This study is based on the establishment of a One Health STEC genomes database, including sequences from isolates of different sources. Most of the isolates had been isolated in the ten-years’ time span 2010-2020, in 11 different countries, for surveillance and monitoring activities or specific surveys and research purposes. The final dataset included the whole genome sequencing of 3,418 STEC isolates, mainly from human cases of infections. The metadata included the host symptoms, where available, for human STEC strains and the animal source the strains had been isolated from. We set up a pipeline for the harmonized analysis of STEC WGS, called Discover, made available though ARIES webserver or GitHub. The analysis allowed a deep characterization of STEC strains circulating in Europe. We used this resource to assess STEC population structure and to investigate potential associations between genomic features, host reservoirs, and various symptoms associated with STEC infection by PCoA. This analysis highlighted the presence of subpopulation of human STEC associated with specific features. We provide new information useful for risk characterization, as well as a large dataset genome database and associated metadata compiled from STEC strains, representing a valuable resource for the scientific community, enabling further investigations into STEC diversity, evolution, source attribution and public health relevance.

**Data summary:** The authors confirm all supporting data, including sequence data accession numbers, code and protocols have been provided within the article or through supplementary data files. One supplementary method and five supplementary tables are available with the online version of this article

## Introduction

Shiga toxin-producing *Escherichia coli* (STEC) are an important cause of foodborne disease worldwide. Human STEC infections can result in a wide spectrum of symptoms, ranging from mild to bloody diarrhea, haemolytic uremic syndrome (HUS) and potentially death. STEC are defined as *E. coli* strains carrying the Shiga toxin (*stx*) genes, harboured on lysogenic bacteriophages, encoding potent cytotoxins able to block protein synthesis in target cells (1). STEC also harbour various additional factors encoded by genes acquired through horizontal gene transfer (2). This leads to substantial variability within STEC populations, resulting in multiple subpopulations characterized by distinct arrays of virulence genes. These gene profiles have been differentially associated with several symptoms observed in human disease (3).

Two main types of Shiga toxins (Stx) have been described, Stx1 and Stx2, of which different subtypes can be recognized (3–6). The subtypes most often associated with the more severe forms of disease and outbreaks are those encoded by *stx2a*, *stx2c* and *stx2d* genes (1,7).

Besides STEC, the diverse group of pathogenic *E. coli* also includes other *E. coli* strains that can cause a wide range of enteric diseases (diarrheagenic *E. coli*, DEC) and systemic diseases (Extraintestinal Pathogenic *E. coli*, ExPEC) in both humans and animals (8,9). However, STEC is the only pathotype with a well-established zoonotic origin. Additionally, pathogenic *E. coli* strains possessing virulence traits of multiple pathotypes have been reported (10,11).

Human STEC infections are frequently reported in the European Union (EU): in 2024, STEC was the third most reported zoonosis in humans, with 11,738 confirmed cases (12). STEC can be isolated from different sources and a variety of food-producing animals, as well as wildlife. Domestic ruminants (e.g. cattle, goats, sheep) are frequently healthy carriers of STEC and have been identified as the primary reservoir of human STEC infections (13–15). However, foodstuffs of ruminant origin, such as hamburgers, minced meat, undercooked beef, as well as unpasteurized milk and cheeses, are seldom identified as vehicles of transmission of STEC to humans. Sources of STEC infection beyond animal-derived food, such as contact with STEC-colonized animals or their environments, wildlife, vegetables, and recreational and drinking water, have been increasingly reported (16,17).

Besides the presence of *stx* genes, STEC possess accessory virulence features carried by mobile genetic elements, acquired through horizontal gene transfer, such as pathogenicity islands (PAIs) and plasmids. This is the case of the intimin adhesin, responsible for the adhesion mechanism known as “attaching and effacing”. Intimin is encoded by the *eae* gene present on the PAI termed Locus for Enterocyte Effacement (LEE) and it is a typical feature of certain STEC frequently isolated form cases of severe disease (18).

Historically, the seropathotype classification of STEC has been applied to account for the diversity of disease risk outcomes (19). However, the serogroup alone does neither account for virulence of STEC strains nor represent an efficient epidemiological marker to track the disease or geographical risk associated with STECs. Typing specific STEC strain characteristics, such as virulence gene determination and *stx* gene subtyping (20), may help identifying pathogenic strains and the associated vehicles of human infection. The application of molecular-based typing methods offers an opportunity to refine risk assessment. In recent years, advanced molecular biology techniques have emerged, ranging from microarray and high-throughput PCR systems targeting virulence profiles to whole genome sequencing (WGS). WGS offers at the moment the greatest potential for enhancing and supporting microbial risk assessment, as the data obtained provides valuable information on strain characteristics at once, including serotype, virulence and antibiotic resistance profiles, as well as enabling genomic typing through core-genome Multi-Locus Sequence Typing (cgMLST) analysis and/or Single Nucleotide Polymorphisms (SNPs), to assess strains relatedness (21).

The DiSCoVeR project (“Discovering the sources of *Salmonella*, *Campylobacter*, STEC and antimicrobial resistance”, https://onehealthejp.eu/jrp-discover/), co-funded by the One Health European Joint Programme, aimed to address knowledge gaps regarding potential sources of human STEC infections. An inventory of STEC isolates from human cases and different animal, food and environmental sources was set up by 11 DiSCoVeR consortium partners in ten EU/EEA countries and UK to perform comparative genomics and source attribution analyses. The inventory included 7,549 STEC isolates from the consortium partners’ collections, mainly obtained through surveillance and research activities in 2010-2020, with data on the basic virulotype (presence of *stx* and *eae* genes) and some information concerning the source. This dataset has been used to carry out a source attribution analysis (D-JRPFBZ-1-WP4.3 available at https://zenodo.org/records/7406419). A subset of these STEC strains has been typed using WGS for a high-resolution characterization of the STEC strains available in the DiSCoVeR database to carry out further genomic analyses.

We present here the characterization of this STEC dataset by WGS, along with subsequent analyses aimed to assess STEC population structure and to investigate potential associations between genomic features, host reservoirs, and the range of symptoms associated with STEC infection.

## Methods

### STEC strains

In total, 3,568 STEC isolates from eleven European countries were sequenced and the related WGS data have been included in this study. The majority of the sequenced isolates (97.2%) were collected between 2010 and 2020, with a smaller number isolated before 2010, and a few in 2021. Isolates and isolation methods have been described elsewhere (https://zenodo.org/records/7406419). The complete cleaned dataset concerning the STEC strains included in this study and the information on the year of isolation, as well as the source, can be found in the supplementary material (Supplementary Table S1, file Supplementary_table_S1_STEC_Discover).

### DNA extraction and whole genome sequencing

DNA extraction, quality and quantity check of DNA, as well as library preparation and sequencing were performed by each partner. Several consortium partners perform sequencing on a routine basis and the sequencing methods used varied according to the standard operating procedures in place at each laboratory. Most isolates were sequenced using enzymatic fragmentation, followed by sequencing paired end reads of 150 or 300 bp using Illumina technology. Several isolates were also sequenced by IonTorrent technology.

### Bioinformatics

A bioinformatics workflow named “Discover” was developed to standardize WGS analyses of STEC strains. The pipeline, written in Python 3.7, was implemented in the publicly accessible Galaxy instance ARIES (https://aries.iss.it) and is also available on GitHub (https://github.com/lucadesabato/Discover1.1). ARIES provides a user-friendly environment, enabling both expert and non-specialist users to analyze raw sequencing data in a harmonized manner.

The pipeline accepts Illumina or Ion Torrent FASTQ files (including compressed versions,.gz) as input. Reads were quality-trimmed with Trimmomatic (Bolger et al., 2014) and assembled using SPAdes v3.15.2 (Bankevich et al., 2012) with subsequent filtering of low-quality scaffolds. Virulence genes were identified using Abricate against the Statens Serum Institut (SSI) database (Joensen et al., 2014), while subtyping of Shiga toxin genes employed the Shiga toxin-typer tool, combining mapping, assembly, and BLAST comparisons to the STSTDB database (Joensen et al., 2014). Serotyping (O:H) was performed with the *E. coli* Serotyper tool (Joensen et al., 2015), and MLST was determined with the mlst software (https://github.com/tseemann/mlst). Antimicrobial resistance genes were detected using Abricate with the ResFinder database (Zankari et al., 2012). Core genome MLST (cgMLST) analyses were conducted using chewBBACA with the INNUENDO schema comprising 2,360 loci (Llarena et al., 2018). The cgMLST analysis was also used as a final checkpoint for the quality of the assemblies. A cut-off for accepting an isolate’s genome was set at ≥ 80% of loci covered (EXC+INF), (≥ 1,888 loci identified and covered).

The main steps of Discover pipeline with software and parameters are listed in Supplementary Method S1.

Assembly quality was evaluated using CheckM (Parks et al., 2015) and QUAST (Gurevich et al., 2013). Cut-offs included >95% completeness, <5% contamination, ≤500 contigs, N50 ≥15,000 bp, and genome size (∼5 Mbp, 50% GC content). A cgMLST coverage threshold of ≥80% (≥1,888 loci) was applied as a final quality criterion.

Results were compiled into structured tables summarizing genome statistics, serotype, MLST and cgMLST profiles, virulence and antimicrobial resistance gene content, and contamination checks. The table merging the results of WGS analysis of all STEC strains included in this study is available as Supplementary Table S1 (file nameSupplementary_table_S1_STEC_Discover).

### Principal Component Analysis (PCoA) and statistical analysis

Data from the DiSCoVeR bioinformatics pipeline were included in the PCoA. The data, containing the presence and absence of virulence genes, were first separated between human and non-human bacterial isolates. From the dataset containing non-human isolates two sets were created; the *stx* genes were removed from both sets, whereas the Locus of enterocyte effacement (LEE) genes (*eae*, *espA*, *espB*, *espF* and *tir*) were removed from only one set. The virulence genes were then one-hot encoded, meaning that for each isolate the presence or absence of each gene subtype was recorded. This was done to fit the data into the PCoA. A 5-component PCoA was performed using the PCoA module from the scikit-learn package for python (22). From the PCoA analyses, we extracted the features, the explained variance, the explained variance ratio and the component information to assess the relevance of PCoA components and to evaluate the importance of different virulence genes for the separation of data. From the databases of the PCoA analyses, all virulence genes with a statistical significance of p<0.001 were extracted using an if-loop in R (https://www.r-project.org/). Subsequently, the significant genetic variants were searched for aggregation using the dplyr package (doi: 10.32614/CRAN.package.dplyr) in the initial database of all STEC isolates, characterized by qualitative and quantitative variables, to determine whether the presence of certain features significantly explained the variance observed in the populations resulting from the PCoA.

### cgMLST-based clustering tree

A minimum spanning tree based on cgMLST allelic profiles was constructed to visualize genetic relatedness among the STEC isolates and was created using the MSTreeV2 method in GrapeTree (23). The tree was visualized using iTOL (24).

### Sequence data availability

WGS data are available at ENA or NCBI and accession number corresponding to each STEC isolate can be found in Supplementary Table S1.

## Results

### Data collection and descriptive statistics of the data set

The database used in this study was composed of genomes and descriptive metadata (Supplementary Table S1) shared by the *DiSCoVeR* project’s consortium partners. Due to the absence of harmonized programs for STEC monitoring and surveillance across the EU, the data provided by each partner reflected strains present in their respective culture-based collections. The consortium partners were requested to provide data without any restriction regarding isolation strategy or sampling plan, with the aim of including as many isolates as possible. The STEC strains included in the database were isolated in different settings (*ad hoc* surveys, monitoring, research projects, surveillance) and with different analytical approaches (e.g. determination of the presence of specific serogroups such as O157). The data collected included genomes of STEC isolates from human and non-human sources from eleven European countries. Five countries, namely Italy, the Netherlands, Norway, Portugal and the United Kingdom, shared genomic data from both human and non-human isolates. Denmark only contributed data from human isolates, while the Czech Republic, Ireland, Poland, Spain and Sweden only shared WGS of strains isolated from non-human sources. Most of the human strains had been isolated in the period 2010-2020 while only a few isolates were from 2003 and 2021. The non-human isolates were collected between 1998 and 2021 and originated from research projects, including STEC strains isolated in the framework of the *DiSCoVeR* project itself, other projects, outbreak investigations, surveys and monitoring and control programs. Summary tables with the number of isolates per country, year, organization and sources of isolation, as well as details of source subgroup, disease outcome, MLST, serotype, presence of *stx* and *eae,* can be found in Supplementary Table S2.

The initial dataset included WGS data from 3,568 STEC isolates, which were submitted to a stepwise filtering based on several quality parameters (see Supplementary Table S3). This quality check process led to a final dataset consisting of 3,418 STEC isolates, which were divided into 2,405 isolates from human cases and 1,013 isolates from non-human sources. The bar graph in Figure 1 illustrates the distribution of STEC isolation source vs the country of origin. The detailed information, including a few descriptive data, as well as the full WGS characterization obtained through the Discover Pipeline, are reported in the Supplementary Table S1. Of the 2,405 genomes from humans, most of them were provided by Denmark, the Netherlands, Norway, Italy and the UK (Figure 1).

**Figure 1.**
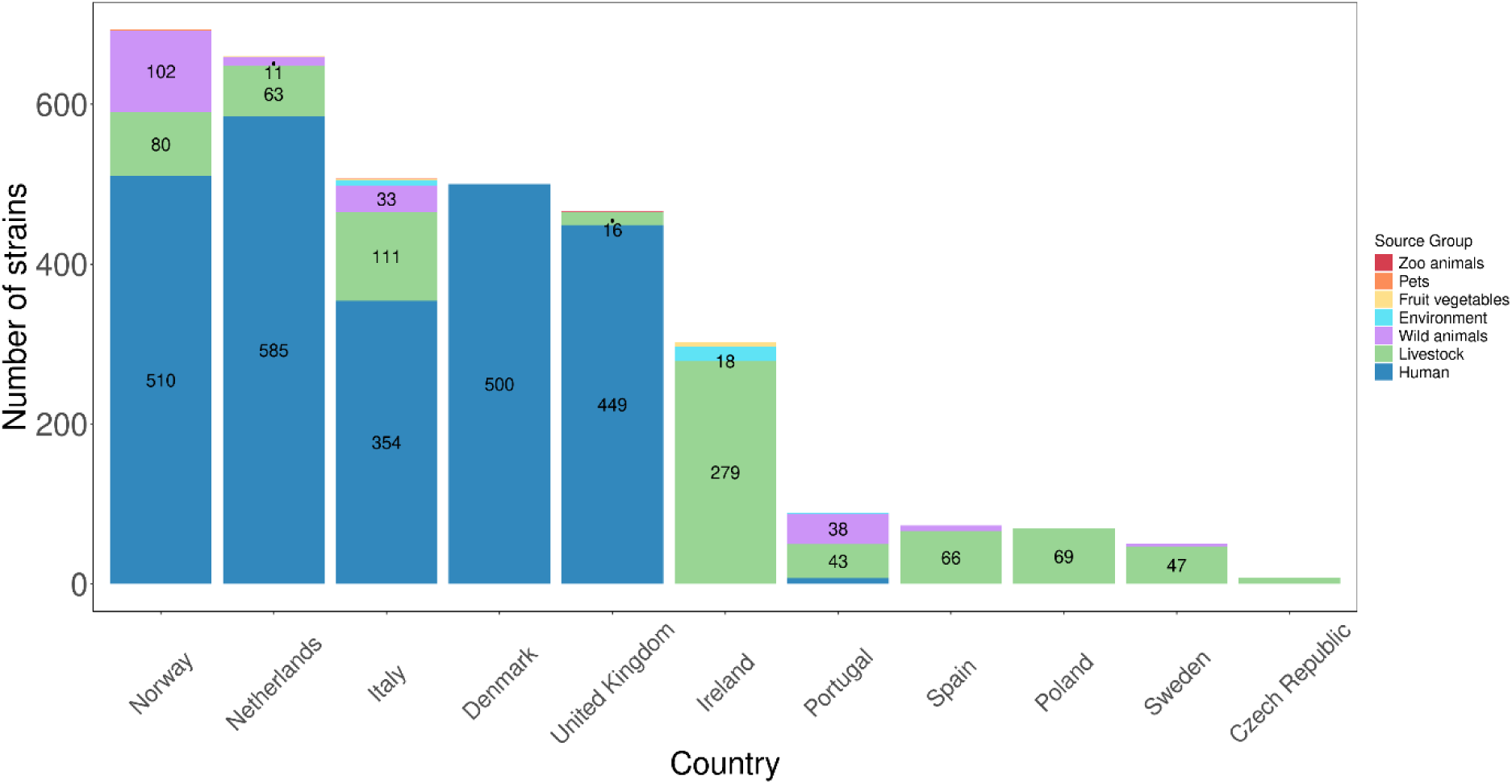
STEC isolates distribution over countries and isolation sources (Source groups). Numbers below ten are not displayed in the figure (Norway: Pets =1; Netherlands: Fruit/vegetables=1; Italy: Fruit/vegetables=1, Pets=1; UK: Zoo animals=1; Ireland: Fruit/vegetables=5; Portugal: Environment=1, Human=7; Spain: Wild animals=7; Sweden: Wild animals=4; Czech Republic: Livestock=8).

As for the human data, the information on the disease status was available for 1,096 genomes out of the total 2,405 human records shared by the consortium partners (Figure 2). Some of the consortium partners could not share the disease outcome from patients due to national regulations and others had not recorded disease outcome associated with all STEC isolates.

**Figure 2.**
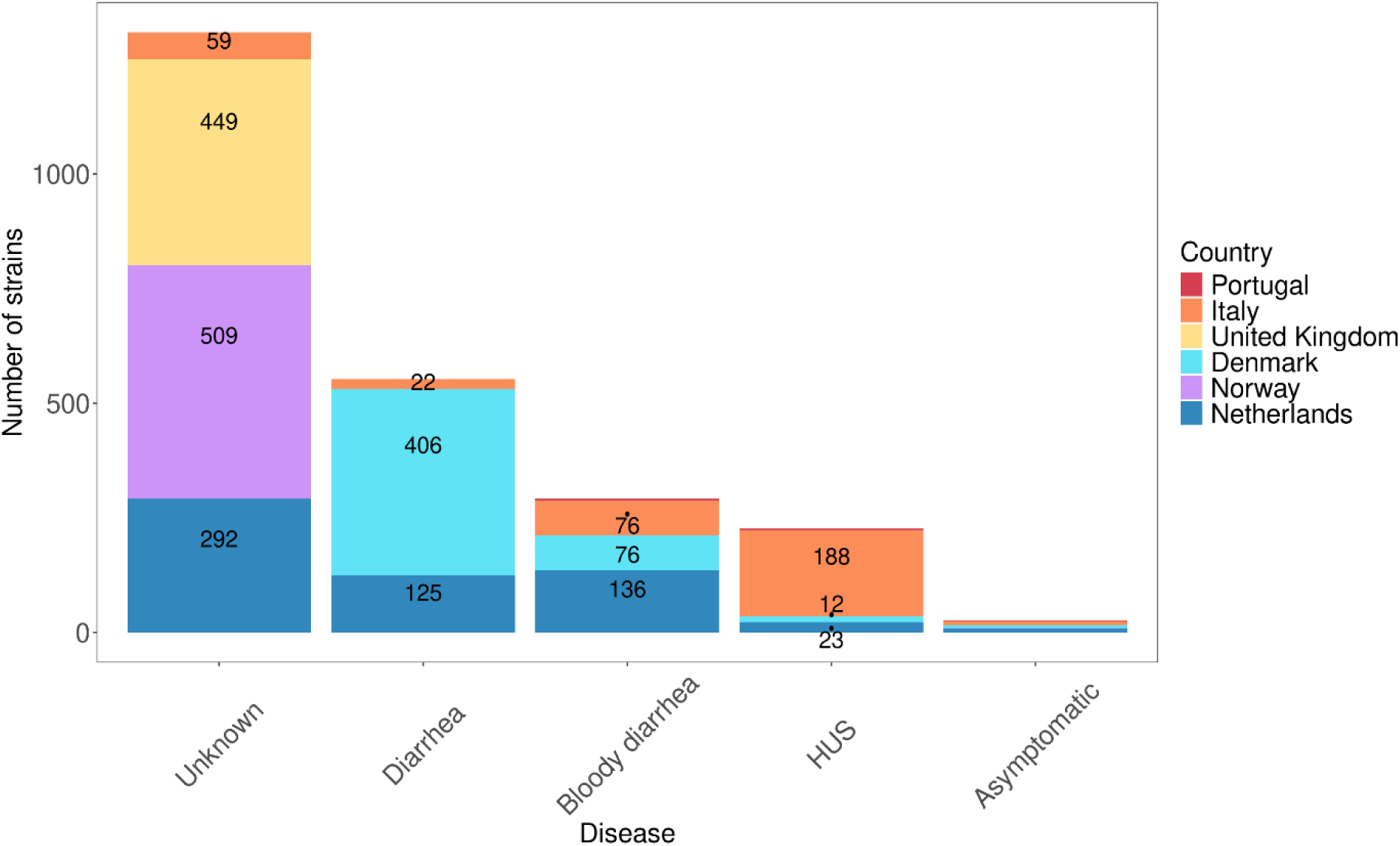
Distribution of STEC strains from human cases based on disease symptoms, by country. Numbers below ten are not displayed in the figure: Portugal=3; Bloody diarrhea: Portugal=3; Asymptomatic: Denmark=6, Italy=9, Netherlands=9, Norway=1, Portugal=1).

As far as the genomes from non-human sources are concerned, each main source group was divided into several subgroups (Figure 3). Genomes of strains from livestock were the predominant source of non-human isolate genomes (n= 782), which included STEC either from live animals, obtained by assaying fecal samples, or food samples like meat and dairy products, followed by wild animals (n= 195). The number of STEC genomes from other source groups such as environment, fruit and vegetables, pets and zoo animals were sparse and only included 36 STEC genomes in total (Figure 1, see also Table S2).

**Figure 3.**
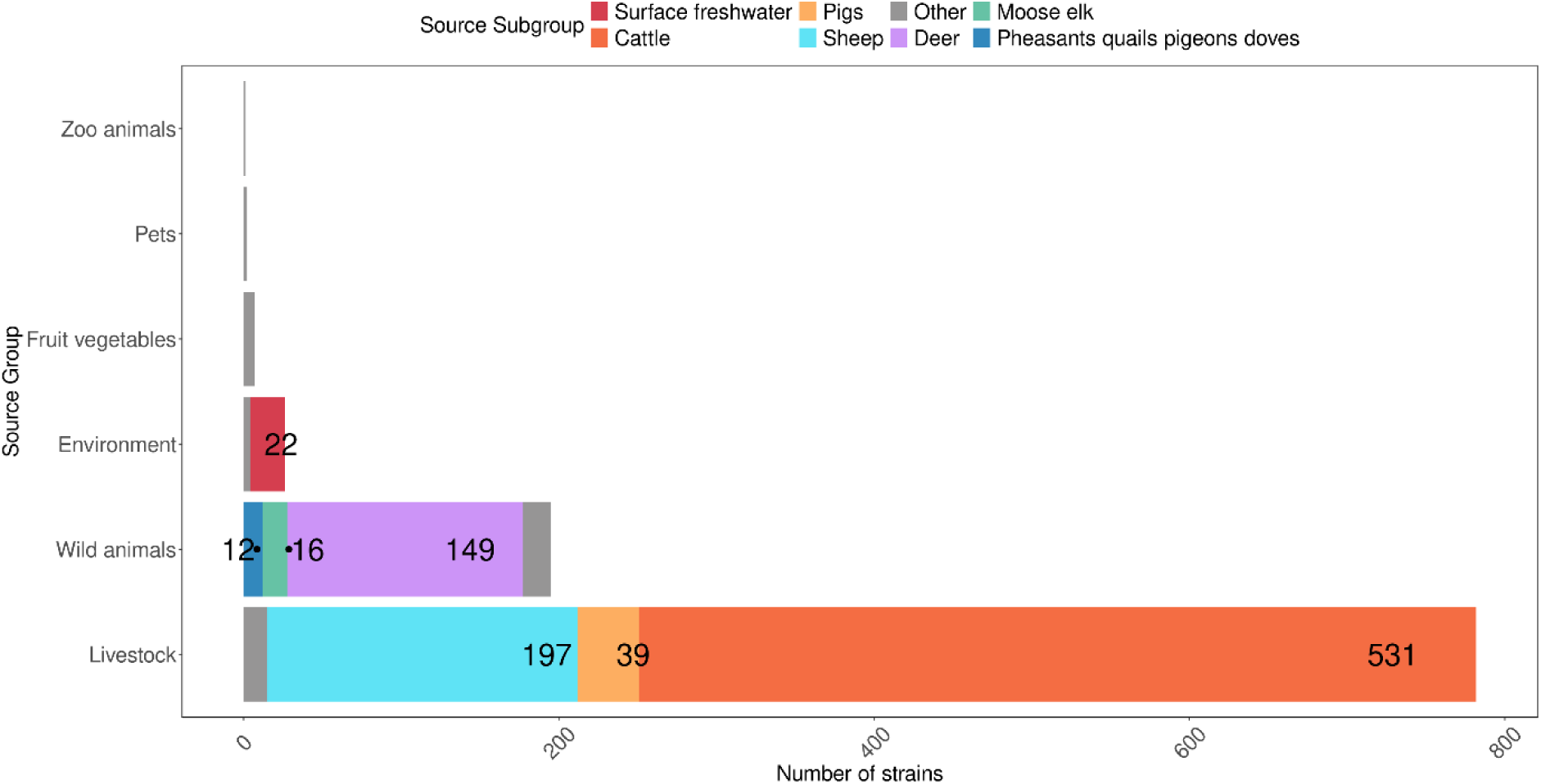
STEC isolates from non-human sources distributed per source group and into different source subgroups. Grey colour represent subgroups from each source group that represent less than ten isolates per subgroup (Zoo animals: Petting goats/sheep=1; Pets: Dogs=2; Fruit/vegetables: Vegetables=6, Nuts/seeds including oils thereof=1; Environment: Groundwater=1, Sewage wastewater=1, Other=2; Wild animals: Ducks, geese and waterfowl=1, Other wild ruminants=8, Rabbits/hares=1, Wild boars=8; Livestock: Farmed shellfish=2, Goats=3, Horses/donkeys=1, Livestock unknown=4, Other mammals=5).

The major source group, livestock, was mainly composed of cattle (n=531), followed by sheep (n=197) and pigs (n=39) (Figure 3). More in detail, of the 782 livestock STEC genomes, 263 were from strains isolated from food, mainly of bovine origin (n=229). The others were from live animals, mainly cattle (n=298) and sheep (n=183).

The wild animal group included genomes from 195 isolates, with most of them (n=147) consisting of isolates from deer (red deer n=77, roe deer n=34, 6 from reindeer, and 30 from unspecified deer species). The wild animal groups also included isolates obtained from moose (n=15) and pigeons (n=12). Only two STEC genomes in this category were from strains isolated from food (deer meat).

### Serotyping and sequence typing

The overview of isolates from human and non-human sources, along with the presence of the main virulence genes and serotype is presented in Figure 4. The basic virulence profile consists in the presence of *stx* gene types and the intimin-coding *eae* gene, considered a hallmark of the LEE locus.

**Figure 4.**
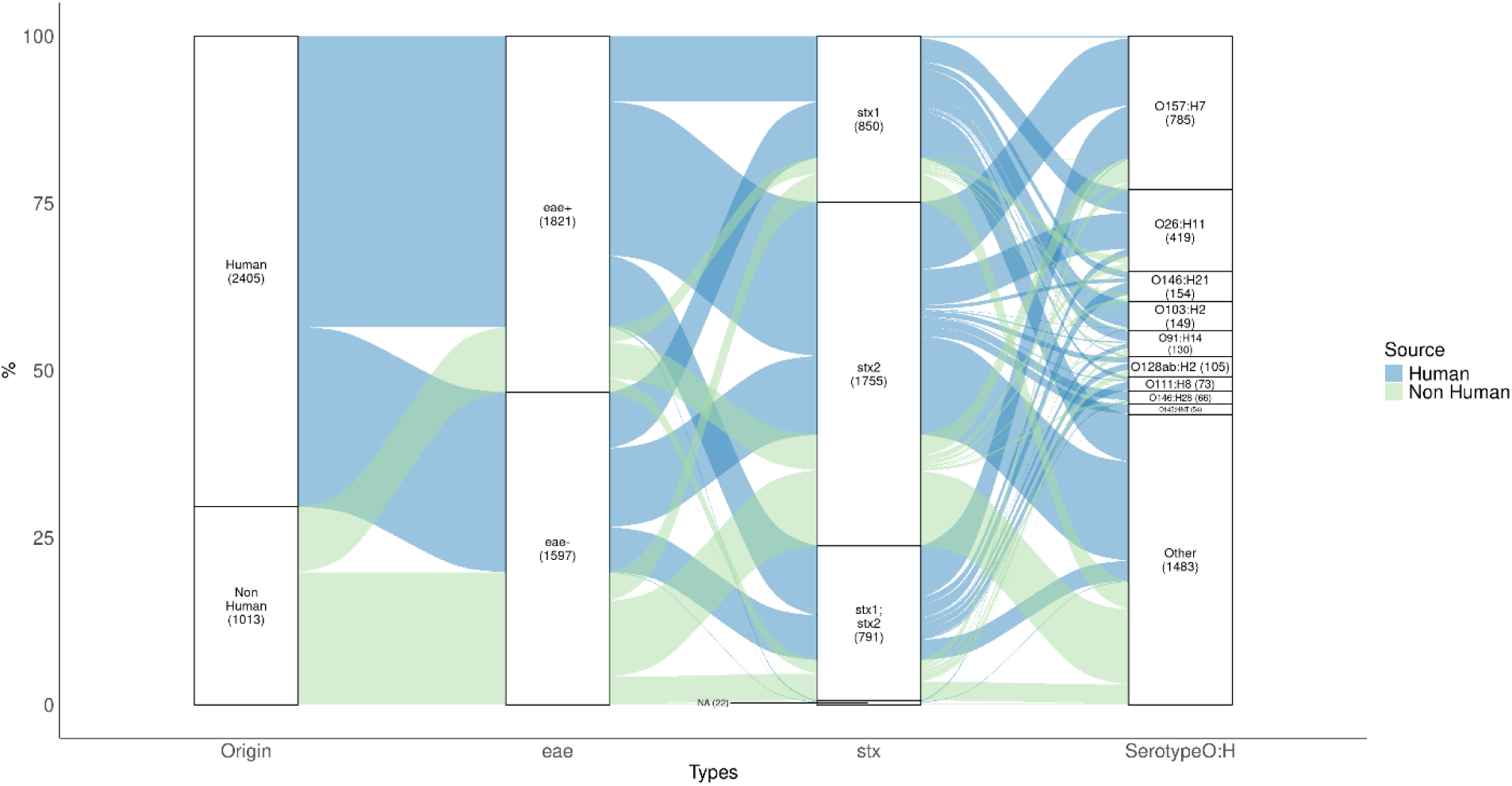
Sankey plot of STEC illustrating different serotypes and virulence genes (*eae* and *stx*) related to human and non-human isolates.

Based on the WGS analysis, the serogroup (O-group) could be determined for 3,131 out of the 3,418 strains, while the H-type could be identified in 3,357 strains. The complete serotyping (determination of both O-group and H-type) was achieved for 3,075 isolates in total (Supplementary Tables S1 and S2).

Overall, 216 different serotypes were identified. The majority of the isolates belonged to serotype O157:H7 (n=785), followed by O26:H11 (n=419), O146:H21 (n=154), O103:H2 (n=149), O91:H14 (n=130), O128ab:H2 (n=105), (Figure 4, Supplementary Tables S1 and S2). The main serotypes of the human STEC isolates with information on disease outcome were O157:H7 (n=242), O26:H11 (n=222), O103:H2 (n=79), O146:H21 (n=27), O91:H14 (n=21) and O128ab:H2 (n=15) (Supplementary Table S1). The figure of the serotype distribution changed when the most severe form of STEC infection, HUS, was considered, with the main serotype being O26:H11 (n=99), followed by O157:H7 (n=48), O111:H8 (n=28), O80:H2 (n=14) and O145:H25/HNT (n=10).

From the non-human STEC isolates, the most common serotypes were O157:H7 (n=155), O26:H11 (n=78), O91:H14 (n=42), O146:H21 (n=37), O128ab:H2 (n=37), followed by O187:H28 (n=30), the latter mainly isolated from wild animals.

Concerning the 7-loci MLST, several sequence types (ST) included more than one serotype. The most frequent sequence types based on MLST was ST11 (n=773) corresponding to O157:H7 and ONT:H7, ST21 (n=343) mainly corresponding to O26:H11, ST17 (n=160) mainly corresponding to O103:H2, ST442 (n=158) mainly corresponding to O146:H2, ST33 (n=131) mainly corresponding to O91:H14, ST10 (n=106) represented in more than 15 serotypes of which O113:H4 being the most common one and ST29 (n=99) mainly corresponding to O26:H11.

### Presence of main virulence genes and cross-pathotypes characteristics

The main virulence genes *stx* were identified in nearly all isolate genomes (n=3396, 99.4%). As all isolates were originally identified as STEC (containing one or several *stx* genes) the lack of *stx* genes in some isolate genomes might be to the result of the loss of *stx*-phages due to long storage and/or (multiple) re-culturing steps or other atypical sequences detected by PCR but not accepted by the strict inclusion criteria for defining presence of a gene in the WGS data.

The *stx1* was identified in 1,641 out of 3,418 STEC genomes, either alone (n=850) or in combination with *stx2* (, while the *stx2* alone was identified in 1,755 isolates (Table 1). The *stx* subtypes in combination with the source are described in Table 1 and per isolate in Supplementary Table S1.

**Table 1:** *stx* genes and *stx* subtypes identified in human and non-human sources.

| ***stx* gene type** | ***stx* subtype** | **Human** | **Non-Human** |
| --- | --- | --- | --- |
| *stx1* | *stx1a* | 482 | 118 |
|  | *stx1c* | 127 | 84 |
|  | *stx1d* | 13 | 26 |
| ***stx1*** | **Total** | **622** | **228** |
| *stx2* | *stx2a* | 499 | 139 |
|  | *stx2b* | 244 | 116 |
|  | *stx2c* | 136 | 112 |
|  | *stx2d* | 45 | 42 |
|  | *stx2e* | 36 | 56 |
|  | *stx2f* | 98 | 15 |
|  | *stx2g* | 17 | 43 |
|  | *stx2a;stx2c* | 91 | 10 |
|  | *stx2a;stx2c;stx2d* | 10 | 2 |
|  | *stx2a;stx2b* | 1 | 12 |
|  | Other | 10 | 21 |
| ***stx2*** | **Total** | **1187** | **568** |
| *stx1;stx2* | *stx1a;stx2a* | 140 | 52 |
|  | *stx1a;stx2b* | 42 | 19 |
|  | *stx1a;stx2c* | 219 | 44 |
|  | *stx1a;stx2d* | 10 | 7 |
|  | *stx1c;stx2b* | 149 | 78 |
|  | Other | 24 | 7 |
| ***stx1;stx2*** | **Total** | **584** | **207** |
| Not Detected | Not applicable | 12 | 10 |

The *eae* gene was identified in more than half of the isolates’ genomes (n=1,821), most of them from humans (n=1,485) (Figure 4).

Analyses of the genomes with the Discover pipeline identified 101 STEC strains harboring the *sta1* gene encoding the heat-stable enterotoxin characteristic of ETEC pathotype. Nineteen of these strains were from humans and possessed *stx2a* (n=2), *stx2e* (n=2) or the *stx2g* gene (n=15). Out of the 82 isolates from non-human sources, 33 were from wild animals, all possessing the *stx2g* subtype (one isolate also contained *stx2b*). Most of the isolates from livestock, as well as the few isolates from fruit/vegetables and one from the environment presenting the *sta1* gene, possessed the *stx2a* subtype, in one case together with *stx1a*. In the database two strains, isolated from humans but with no information on the disease status, possessed the gene encoding the heat labile enterotoxin (*ltcA*), produced by some ETEC strains.

One outbreak related STEC strain from a HUS case (ED0924) that was negative for the *eae* gene but possessed the *stx2a* subtype, showed the genetic features of enteroaggregative *E. coli*, showing the presence of *aggR*, *agg3*, *aat* and *aaiC* genes, among others (Supplementary Table S1).

A more detailed analysis of the virulence genes is described in association with the source of genomes in the PCoA analysis.

### cgMLST analysis

A cluster analysis based on cgMLST was performed on all STEC genomes. The resulting tree is shown in Figure 5. The labels indicate the presence/absence of LEE genes (i.e. *eae*, *espA*, *espB*, *espF*, *tir*), as well as the source from which the isolates were obtained. In general, LEE-positive and LEE-negative STEC cluster separately, suggesting distinct evolutionary origins. Isolates from different sources are dispersed throughout the tree; however, due to the large number of genomes included in this analysis, drawing further conclusions is challenging.

**Figure 5.**
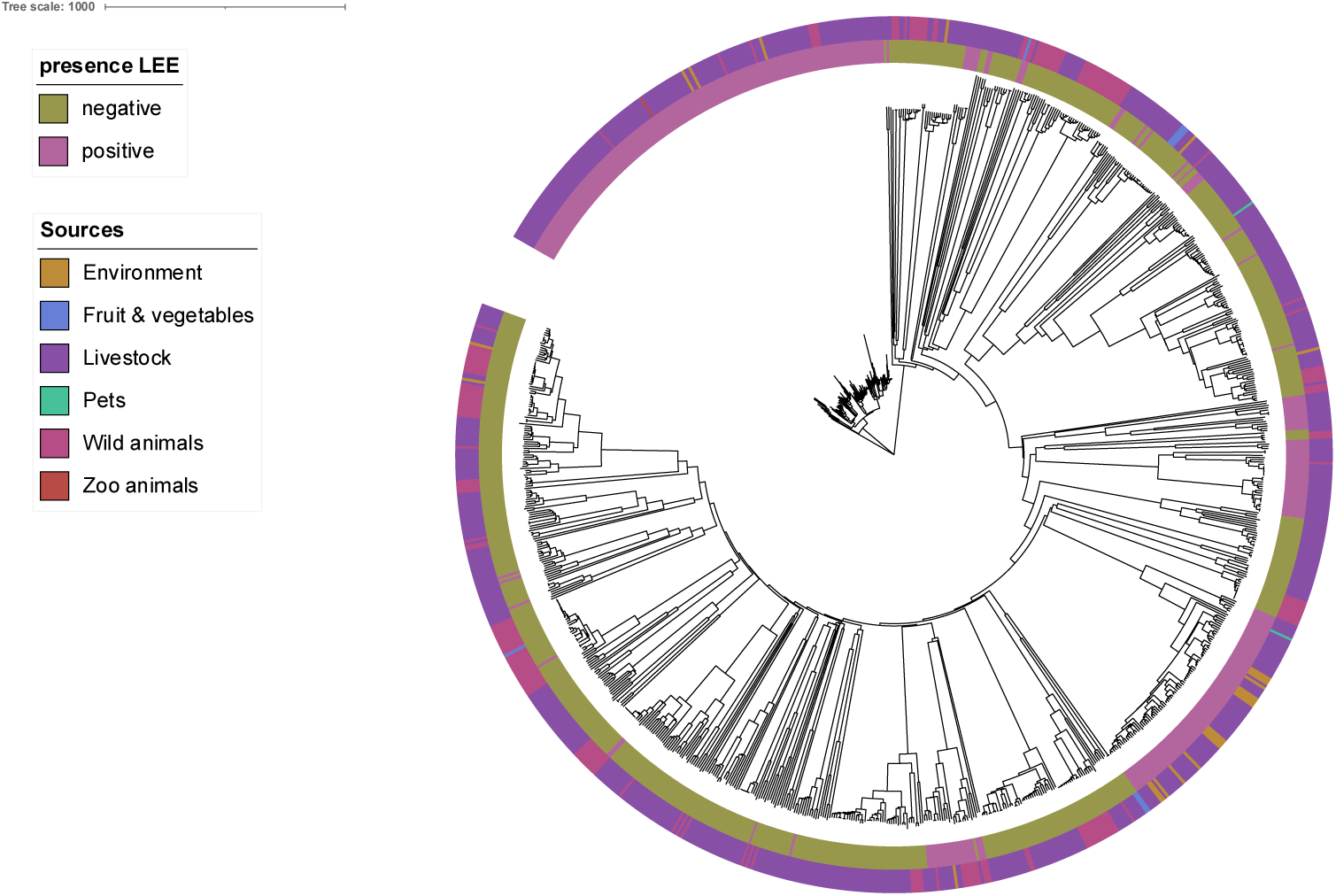
Minimum spanning tree based on cgMLST allelic profiles constructed to visualize genetic relatedness among STEC isolates included in this study. The tree was created using the MSTreeV2 method in GrapeTree and visualized using iTOL.

### Principal component analysis

Principal Components Analysis (PCoA) was applied to the whole dataset to explore potential clustering of specific STEC subpopulations and to identify features explaining the observed variance.

The PCoA analysis revealed three distinct subpopulations among human STEC genomes grouped by symptoms (Figure 6). In particular, the analysis highlighted distinct subpopulations when certain components were considered.

**Figure 6.**
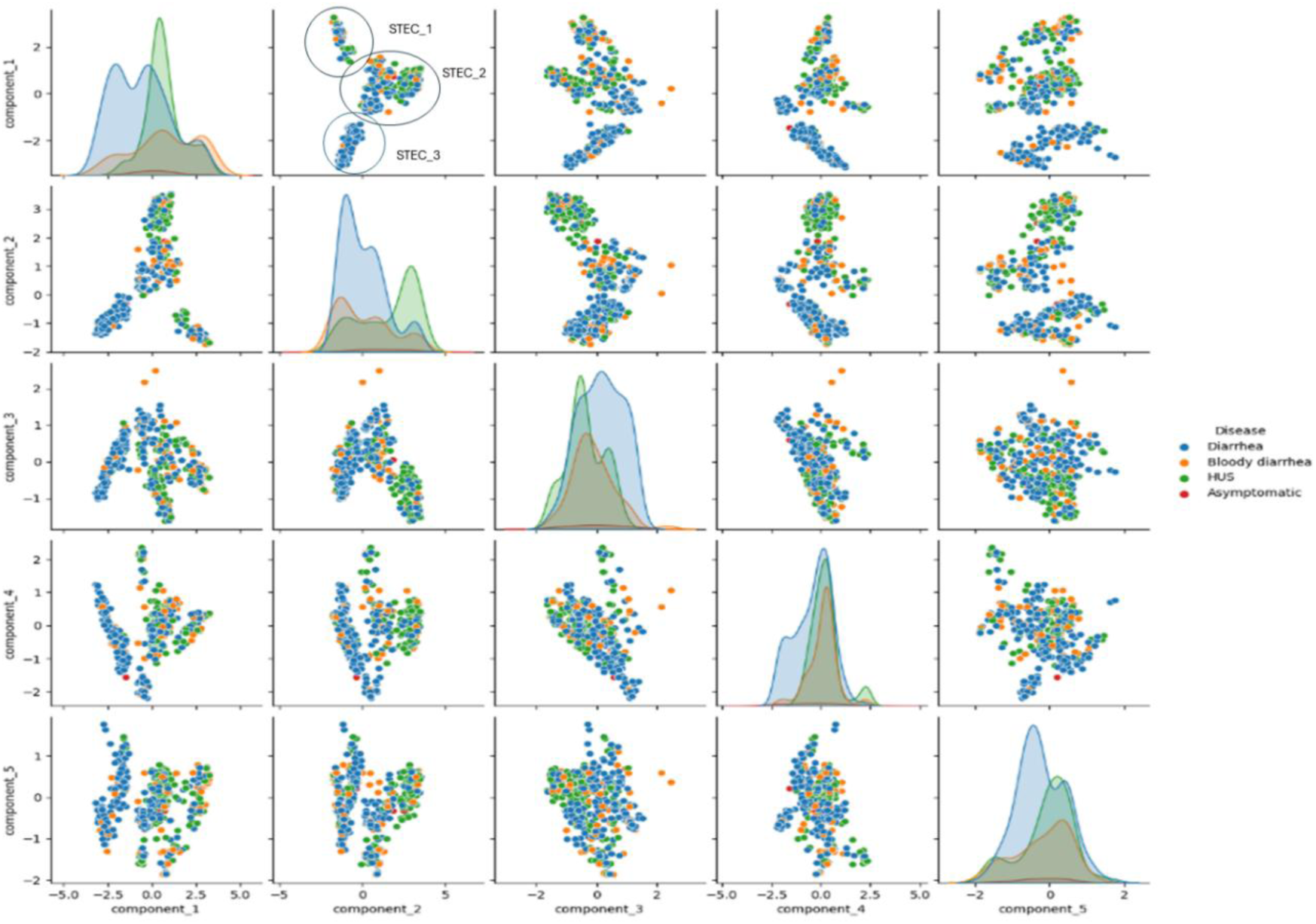
PCoA plot of the genomes from human STEC isolates. The percentage of the variance explained by the single components was: PC1 26.1%; PC2 16.0%; PC3 6.2%; PC4 4.4%; PC5 3.0%.

In addition, to assess whether specific genomic features explained the observed variance, all virulence genes with a statistical significance of p<0.001 were extracted from the PCoA analysis. These genes were searched for aggregation in the initial database of all STEC genomes, to determine whether the presence of certain features significantly explained the variance observed in the STEC_1, STEC_2 and STEC_3 populations identified in the PCoA.

The populations STEC_1 and STEC_3 could be explained by the presence of specific gene patterns (P<0.001).

The identification of the STEC_1 population (286 isolates, Supplementary Table S4) was driven by the presence of microcin-related genes (*mchB, mchC, mchF and mcmA*) linked to intestinal colonization. Additional genes whose presence seemed to explain the variance of the STEC_1 population were siderophore genes (*ireA* and *iroN*) and other the virulence genes (*iss, lpfA*) and*, subA* commonly associated with ExPEC, particularly Uropathogenic UroPathogenic *E. coli* (UPEC) and LEE-negative STEC strains. As a matter of fact, STEC_1 population lacked *eae* gene, encoding the intimin, hallmark of the LEE-positive STEC (Figure 7). Accordingly, 97.9% of these isolates were LEE-negative STEC (Supplementary Table S4), and belonged to 55 different O-groups. Of these, 53.1% were attributed to well-known LEE-negative O groups such as O91, O113, O174 and O146, described in human disease. Table 2 shows the *stx* subtypes associated with disease status in STEC_1. Among 100 isolates with known disease status, 69% were from diarrhea cases, 17% from bloody diarrhea and 12% from HUS (Figure 8). No significant association was found between *stx* subtype and disease status, though *stx1a* and *stx2b* were prevalent in diarrhea cases (Table 2).

**Figure 7.**
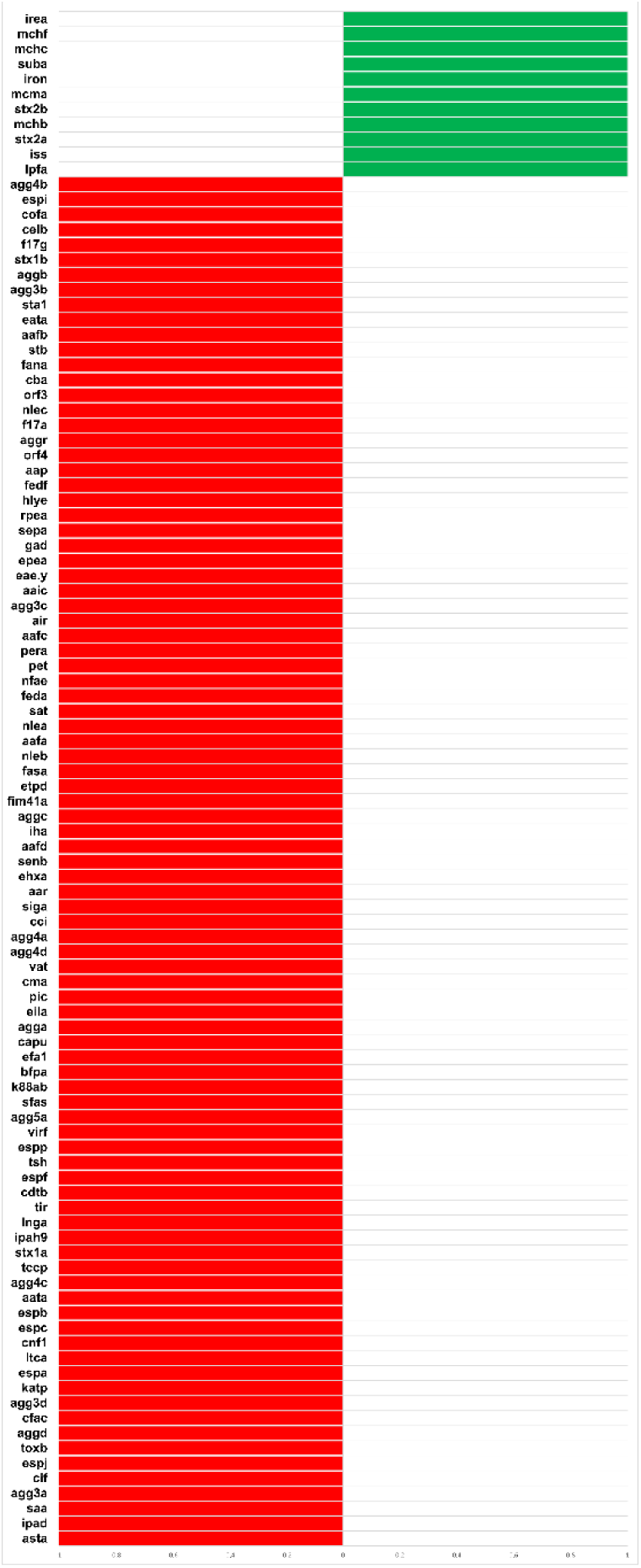
Positive (green) and negative (red) association of the variables within STEC_1 population. In this figure stx2a represents the gene encoding the A subunit of Stx and stx1b and stx2b the genes coding for the B subunit, nor the *stx* genes subtypes.

**Figure 8.**
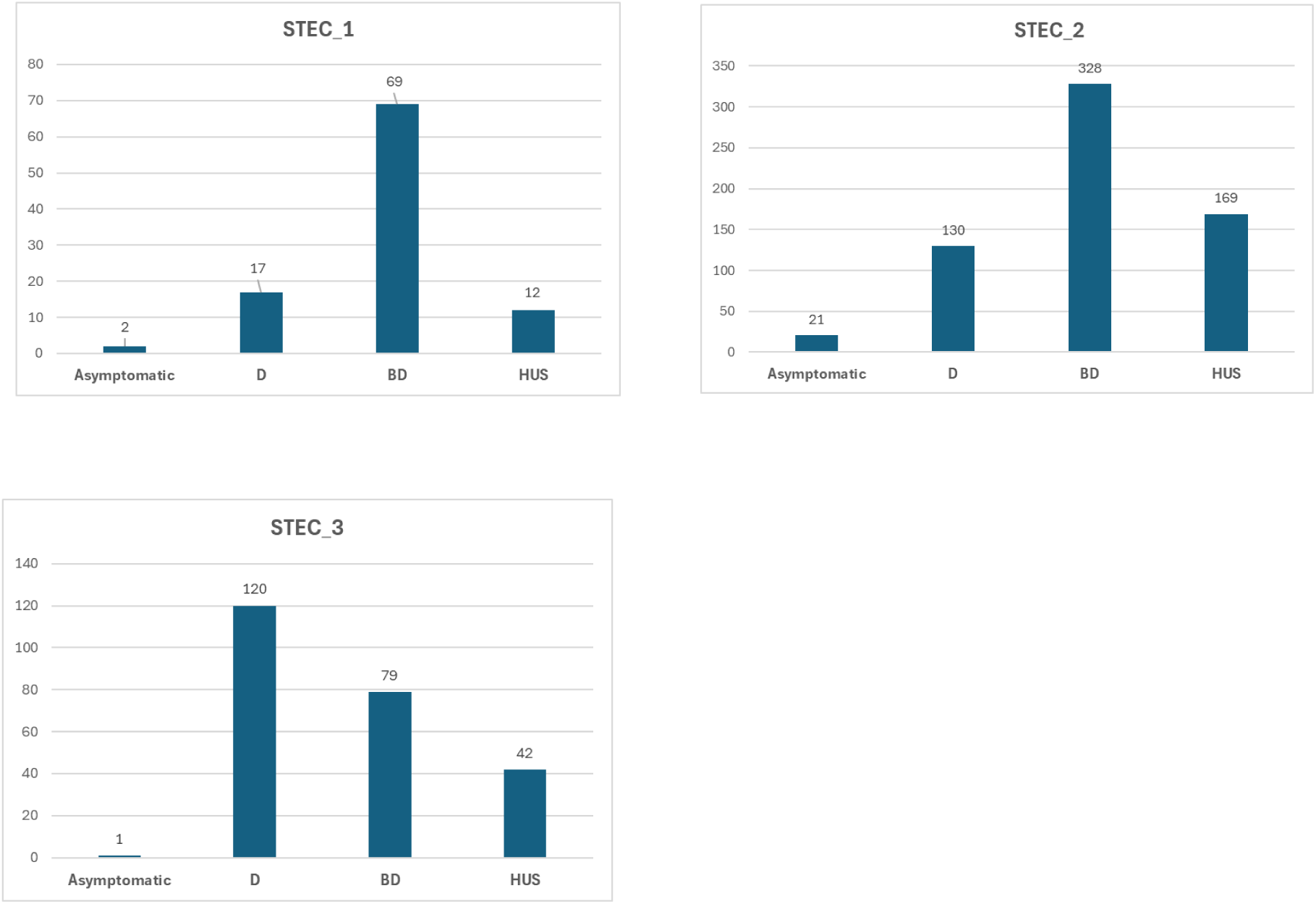
Distribution of the genomes composing the three STEC populations by symptoms. The bar diagrams include: STEC_1 N=100 isolates, STEC_2 N=648, STEC_3 N=242. The genomes from unknown disease status samples are excluded. BD= Bloody Diarrhea, D= Diarrhea, HUS= Haemolytic Uremic Syndrome

**Table 2.** *stx* gene subtypes identified in the genomes of STEC_1 group by the different disease status.

| Diarrhea |  | Bloody diarrhea |  | Haemolytic Uremic Sindrome |  |
| --- | --- | --- | --- | --- | --- |
| *stx* subtype | No. | *stx* subtype | No. | *stx* subtype | No. |
| *stx1a* | 12 | *stx1a* | 3 | *stx1a; stx2a* | 2 |
| *stx1a; stx2a* | 1 | *stx1a; stx2a* | 2 | *stx1a; stx2a; stx2d* | 1 |
| *stx1a; stx2a; stx2c; stx2d* | 1 | *stx1c; stx2b* | 1 | *stx1c; stx2b* | 2 |
| *stx1c* | 3 | *stx1d* | 1 | *stx2a* | 3 |
| *stx1c; stx2b* | 2 | *stx2a* | 1 | *stx2a; stx2d* | 2 |
| *stx1d* | 4 | *stx2a; stx2c* | 1 | *stx2c* | 1 |
| *stx2a* | 3 | *stx2b* | 4 | *stx2d* | 1 |
| *stx2a; stx2c; stx2d* | 1 | *stx2c* | 2 |  |  |
| *stx2a; stx2d* | 1 | *stx2d* | 1 |  |  |
| *stx2b* | 21 | *stx2g* | 1 |  |  |
| *stx2c* | 1 |  |  |  |  |
| *stx2d* | 6 |  |  |  |  |
| *stx2e* | 3 |  |  |  |  |
| *stx2g* | 10 |  |  |  |  |
| **Total** | **69** |  | **17** |  | **12** |

The STEC_2 population (1,263 isolates) included both LEE-positive and LEE-negative STEC across multiple serogroups, notably the well-known O26, O111, O103, O145 as well as O80 of LEE-positive STEC. No significant association of specific virulence gene patterns were identified. In total, 648 isolates had known disease status (Figure 9) and the distribution of *stx* subtypes was much more dispersed in this population than in STEC_1, although the *stx2a* subtype was dominant (73.4%), especially among O26 isolates (Supplementary Table S1). It has to be noted, however, that this serogroup accounted for 58.6% of the isolates from HUS cases, partially explaining the higher proportion of HUS-associated strains in this group and the skewed distribution of the *stx* subtypes towards *stx2a*.

**Figure 9.**
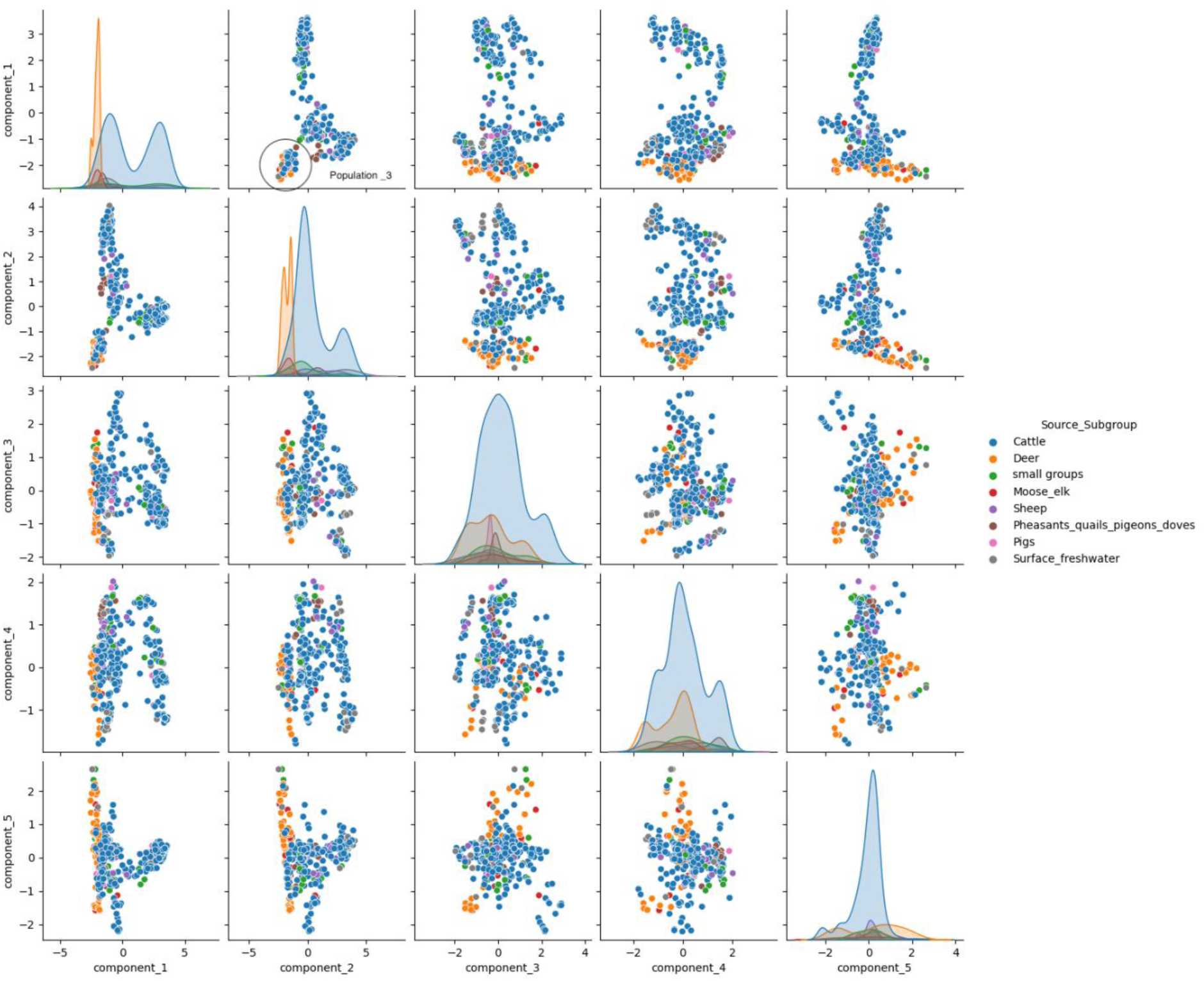
PCoA plot of the genomes from non-human STEC isolates. The percentage of the variance explained by the single components was: PC1 25.6%; PC2 16.5%; PC3 5.9%; PC4 4.4%; PC5 3.9%.

The STEC_3 population (630 isolates) was composed of only STEC O157:H7 (Supplementary Table S5). In total, 242 isolated had known disease status with 32.6% were from diarrhea, 49.6% bloody diarrhea and 17.4% from HUS (Figure 8). No significant association was found between *stx* subtype and disease status. However, the combination of *stx1a* and *stx2c* was commonly found in isolates from diarrhea and bloody diarrhea, respectively, while *stx2a* alone was more frequent in isolates from HUS cases (Table 3).

**Table 3.** *stx* gene subtypes identified in the genomes of STEC_3 group by the different disease status.

| Diarrhea |  | Bloody diarrhea |  | Haemolytic Uremic Syndrome |  |
| --- | --- | --- | --- | --- | --- |
| *stx* subtype | No. | *stx* subtype | No. | *stx* subtype | No. |
| *stx1a* | 1 | *stx1a* | 7 | *stx1a; stx2a* | 7 |
| *stx1a; stx2a* | 17 | *stx1a; stx2a* | 9 | *stx1a; stx2a; stx2c* | 1 |
| *stx1a; stx2c* | 32 | *stx1a; stx2a; stx2c* | 6 | *stx1a; stx2a; stx2c; stx2d* | 2 |
| *stx2a* | 3 | *stx1a; stx2a; stx2c; stx2d* | 1 | *stx1a; stx2c* | 5 |
| *stx2a; stx2c* | 4 | *stx1a; stx2c* | 58 | *stx2a* | 16 |
| *stx2a; stx2c; stx2d* | 1 | *stx2a* | 20 | *stx2a; stx2c* | 5 |
| *stx2c* | 21 | *stx2a; stx2c* | 6 | *stx2c* | 6 |
|  |  | *stx2c* | 13 |  |  |
| **Total** | **79** |  | **120** |  | **42** |

The PCoA analysis of animal STEC isolates revealed a diffuse distribution, with no clear subpopulations except for the one termed as population 3 (highlighted in Figure 9). This group included 122 cattle isolates (23.0%), 78 deer (53.1%), 11 sheep (64.7%) and 25 pig isolates (89.3%), along with smaller groups from other animal sources (Figure 9). Statistical analysis of features associated with this subpopulation showed distinct genes linked to specific host species. Cattle and sheep isolates were significantly associated with LEE locus genes (*eae*, *tir*) and effector genes located outside the LEE region (*nleA*, *nleB*). These isolates also harboured genes found on large STEC virulence plasmids like pO157. STEC isolates from deer displayed features typical of an STEC hybrid cross-pathotype, such as *ireA*, *lpfA*, *mchB*, *mchC*, and *mchF*, also identified in the STEC_1 population. Additionally, *sta1*, encoding the heath-stabile enterotoxin characteristic of ETEC, was present. Pig isolates were characterized by the absence of the LEE locus, the presence of *stx2e* and the *fedF* gene encoding the F18 fimbrial adhesin.

## Discussion

Risk assessment of foodborne pathogens is essential to protect consumers from acquiring foodborne infections, and strain (hazard) characterization is the first step to be undertaken. Knowledge of the pathogen’s population structure, as well as colonization features in animal reservoirs and human pathogenicity is key for source attribution and control strategies. Despite extensive research on STEC diversity and its link to human disease, there is no consensus on prioritizing control measures to reduce infection risk. According to EFSA (2020), all STEC strains are considered pathogenic to humans, capable of causing at least diarrhea. However, strains producing certain Stx subtypes or possessing additional colonization factors may be more prone to cause severe illness, including bloody diarrhea and HUS (20).

To support hazard characterization of STEC, we analyzed a large collection of STEC genomes from different countries collected in the framework of the European project *DiSCoVeR*. Isolates shared from the consortium partners were originally collected for different purposes, including research projects, surveillance activities and *hoc* studies, therefore, they might not be representative of all STEC strains circulating in Europe. The collection included information on STEC strains isolated from different sources (JRPFBZ-DiSCoVeR-1-WP4.3, available at https://zenodo.org/records/7406419). Of these, 3,568 were subjected to WGS for a deeper characterisation and 3,418 were of sufficient quality to be included in this study. The genomic analysis showed that most isolates from human cases with known disease status belonged to the top five serogroups, with O157 representing the highest rank, followed by serogroup O26 (Figure 4). This distribution changed when the severe forms of infections were considered, with O26 becoming the most common serogroup in HUS cases (Figure 4). This is in agreement with numbers reported by ECDC/EFSA in 2024 (12). The most common serogroups identified in non-human isolates were also represented among strains from human disease, displaying similar pattern as reported annually to the EFSA/ECDC (26,27). It has to be noted however, that isolation protocols for STEC may be biased towards isolating these serogroups.

A multidimensional PCoA revealed three distinct human STEC subpopulations (STEC_1, STEC_2 and STEC_3), each associated with different virulence profiles and disease outcome. The STEC_1 subpopulation comprised STEC strains with a hybrid virulence profile, combining *stx* genes with ExPEC-associated traits (28). Such a hybrid *E. coli* pathotype has recently been described as an emerging STEC population causing human disease, including the severe forms bloody diarrhea and HUS (10,29). STEC isolates in the STEC_1 group shared genes encoding microcin production (*mchBCF* and *mcmA*), as well as genes encoding a siderophore system and serum resistance (30–33). A common feature among the genomes in this population was the absence of the LEE locus as well as the large virulence plasmid pO157, typical of STEC O157 and other LEE-positive strains (34,35). This finding confirms that although the LEE locus is a factor favoring the progression of the infection in humans towards the more severe forms, this locus may not be crucial (20). Accordingly, the STEC_1 population included strains isolated from cases of disease characterized by different degrees of severity (Figure 8). Two thirds of the isolates with known disease status originated from diarrheal cases, while approximately the remaining third was from severe cases of bloody diarrhea and HUS (Figure 8).

The STEC_2 group included a wider population of genomes (Figure 6) with both LEE-positive and LEE negative strains. Half of the isolates were from cases of diarrhea, while the others were distributed among the disease status bloody diarrhea and HUS with 20.1% and 26.1% of the genomes, respectively (Figure 8). The STEC_2 group contained the LEE-positive top five STEC serogroups except for the O157 and included the emerging O80 serogroup, which also shows the presence of the LEE locus and of certain traits shared by ExPEC (36). Statistical analysis did not identify any specific genes (presence or absence) describing this population neither to subgroup isolates nor to disease status. This finding suggests that in this case the attribution of the outcome may not be inferred from the strains’ features, but other factors may concur, as it has been hypothesized for the host’s protease profile (37). A limitation of the inference based on the genomic analysis of the STEC_2 population may lie in the overrepresentation of STEC O26 genomes, which accounted for 58.6% of the HUS related entries, thereby skewing the dataset towards a well-known HUS associated STEC serogroup (38).

The STEC_3 population was clearly distinct from the other two (Figure 6). Interestingly, it included 630 genomes from STEC O157:H7. The distribution of isolates with confirmed disease status confirmed that this serotype can cause the full spectrum of symptoms, albeit with varying frequencies. Our dataset showed a peak related to isolates from patients with diarrhea and bloody diarrhea, while only 17.4% of the STEC O157:H7 were associated with HUS cases, as previously observed in this and other studies (20). Our analysis also confirmed the importance of the *stx2a* subtype and its association with strains from HUS, as stated in the EFSA opinion regarding pathogenicity assessment of STEC (20). As in other studies, this subtype has been found in most of the isolates from HUS cases in the populations studied, making it the most reliable feature for proactively identifying the strains potentially involved in causing HUS.

The PCoA applied to the STEC from non-human sources displayed a more dispersed distribution, with no distinct populations except for one (Figure 9, Population 3). This population included STEC isolated from multiple animal species, and the statistical analysis showed significant associations between specific genes sets and the animal source. The LEE-positive STECs were predominantly isolated from cattle and sheep.

These animal species have been described as natural reservoirs of STEC O157, but other LEE positive STEC serogroups are also frequently isolated from their fae samples (13,39). An interesting association was found between certain gene patterns in STEC from deer. These isolates were LEE-negative, but possessed several genes associated with ExPEC, similarly to what was observed in the isolates comprising the STEC_1 population. The STEC from deer encoded the microcin-encoding genes (*mchB*, *mchC,* and *mchF)* and determinants associated with iron metabolism (*ireA* and *lpfA*), suggesting that these species might be a possible reservoir of hybrid STEC-ExPEC strains. The deer population also contains STEC with other cross-pathotype features, such as those containing the gene encoding the heat-stabile enterotoxin of ETEC, *sta1* (9).

We also identified an association between specific genomic features, such as *stx2e* and the *fedF* gene, encoding a fimbrial adhesin (40) and STEC isolates from pigs. The presence of these two genes with this subpopulation is not surprising, as both encode factors produced by *E. coli* strains known to cause disease in pigs. Indeed, *stx2e* is associated with the oedema disease in pigs while F18 fimbriae are associated with ETEC strains, which are a leading cause of postweaning diarrhea (40).

In conclusion, we established a European dataset comprising STEC genomes from various sources across multiple countries, accompanied by a database of epidemiological metadata. We used this resource to assess STEC population structure and to investigate potential associations between genomic features, host reservoirs, and various symptoms associated with STEC infection. Our analysis identified distinct subpopulations of STEC from human cases, each characterized by specific genetic features and associated with varying proportions of severe disease outcomes. Notably, we found that STEC genomes from animal sources displayed genomic patterns associated with specific host species. One key finding was the overlap between the STEC_1 subpopulation—linked to severe human disease—and a group of STEC genomes isolated from deer, highlighting a potential zoonotic link.

We provided novel insights supporting the risk assessment of STEC and described a dataset that can be further used to unravel the circulation of this pathogen along the pathway from the animal reservoir to the human host. Moreover, we identified strains carrying cross-pathotype virulence gene arrangements, comprising genes typical of both STEC and other pathotypes like ExPEC or ETEC. This finding underlines the power of genomic analyses in revealing the full spectrum of virulence determinants in these pathogens, which may be overlooked by assays targeting only limited sets of genes. In addition, the genome database and associated metadata compiled from STEC strains shared by the consortium partners provides a valuable resource for the scientific community, enabling further investigations into STEC diversity, evolution, source attribution and public health relevance.

## Author statements

### Author contributions

The design of the study was conceived by S.M., R.T., C.S. and T.S., data collection, analysis and curation, all authors; the Discover pipeline was deployed by A.K. and L.D.S., formal analysis, T.S., S.M., R.T. and M.M.; writing—original draft preparation, R.T., S.M. and C.S.; writing—review and editing, all. Finally, all the authors revised the manuscript and approved the final version.

## Acknowledgements

The authors gratefully acknowledge the contributions of colleagues at partner institutions for their support in STEC isolation and whole genome sequencing.

Corresponding author would like to thank Manuela Marra and Valentina Libri from the Core Facilities Technical—Scientific Service Team at Istituto Superiore di Sanità for the whole-genome sequencing of the majority of the isolates from Italy.

In Norway, parts of bioinformatic work were performed on resources provided by UNINETT Sigma2, the National Infrastructure for High Performance Computing and Data Storage.

Marie Chattaway is affiliated to the National Institute for Health Research Health Protection Research Unit (NIHR HPRU) in Gastrointestinal Infections at University of Norwich and Public Health Genomics at University of Birmingham in partnership with UKHSA, in collaboration with University of Newcastle, and are based at UKHSA. The views expressed are those of the author(s) and not necessarily those of the NHS, the NIHR, the Department of Health and Social Care or UKHSA.

