## Supplementary_M&M for "Population analysis and host-disease associations of Shiga toxin-producing *Escherichia coli* from various sources across eleven European countries using whole genome sequencing"

**Supplementary Materials and Methods**

**Discover pipeline steps**

The nine steps of Discover pipeline and the related parameters for each software used are listed below:

1) Trimming: The base trimming process of sequencing positions with low Phred values was carried out using the Trimmomatic software (Bolger et al., 2014). The chosen options were: SLIDINGWINDOW, LEADING, TRAILING, MINLEN)

2) Construction of the scaffolds: carried out using SPAdes software, Version 3.15.2 (St. Petersburg genome assembler) (Bankevich et al., 2012). Scaffolds were then filtered using the following options: SpadesCoverage cut-off ratio (0.33), SpadesRepeat cut-off ratio (1.75), SpadesLength cut-off (1,000), Spades coverage-length-cutoff (default = 5,000).

3) Screening of virulence factors: performed on scaffolds using the Abricate software (https://github.com/tseemann/abricate) against the database of virulence genes developed by the Statens Serum Institut (SSI) in Copenhagen (Joensen et al., 2014). The database is available on the website of the Center for Genomic Epidemiology (CGE: http://www.genomicepidemiology.org) (accessed October 2022) and contains 145 virulence genes described in STEC strains, each present in different allelic variants.

4) Subtyping of the genes coding for Shiga toxins: carried out with the Shiga toxin-typer software (https://github.com/aknijn/shigatoxin-galaxy), which took as input the raw data and the scaffolds. The raw reads were aligned to the Shiga toxin references using duk tool. The aligned reads were used to build novel scaffolds using SPAdes software, Version 3.15.2, and SKESA tool. The novel scaffolds, and those built in point 2 were compared by the BLASTN algorithm against the database containing the alleles of the genes coding for Shiga toxins (STSTDB) produced by the SSI and distributed by the Technical University of Denmark (DTU) (Joensen et al., 2014).

5) Serotyping (O:H): carried out with the *E. coli* Serotyper software (<https://aries.iss.it/>) and the database developed by the SSI (Joensen et al., 2015). The database included the genes encoding 188 O antigens and 56 H antigens.

6) Multi-locus Sequence Typing (MLST): performed by the mlst software (https://github.com/tseemann/mlst) with scaffolds as input data. It returned the corresponding MLST schema, Sequence Typing (ST), allele IDs, and allelic variants.

7) Screening for antibiotic resistance genes was carried out by comparing the scaffolds against the Resfinder database (Zankari et al., 2012) implemented in the Abricate tool (2434 nucleotide sequences, 2019-Jul28, https://github.com/ tseemann/abricate/).

8) Core Genome Multi-locus Sequence Typing (cgMLST): The determination of the cgMLST was performed using the chewBBACA software (Comprehensive and Highly Efficient Workflow BSR-Based Allele Calling Algorithm) on assemblies with a predefined schema developed by the INNUENDO project (Llarena et al., 2018), including 2360 loci.

9) Results collection and formatting: the results produced by the Discover pipeline were combined in a tabular format and aggregated into four tables. Each row contained the results of the analysis of a single bacterial genome. The first table includes information on Average Scaffold coverage, cgMLST, MLST, and the housekeeping genes with the reported allelic variant, the subtype of genes encoding Shiga toxins (Stx subtype), serotype (O:H), and the allelic variant of each of the 145 virulence genes. The second table contains the AMR genes found, the third table contains the cgMLST loci and the reported allelic variants, and a fourth table contains information on the species identified and the possible presence of repeated *loci*, for the purpose of assessing possible sequences contamination.
